# Methodological Considerations When Using Polygenic Scores to Explore Parent-Offspring Genetic Nurturing Effects

**DOI:** 10.1101/2023.03.10.532118

**Authors:** M. Chuong, M.J. Adams, A.S.F. Kwong, C.S. Haley, C. Amador, A.M. McIntosh

**Author notes:** joint last authors (C.S. Haley, C. Amador, A.M. McIntosh).

## Abstract

**Background:** Research has begun to explore the effects of parental genetic nurturing on offspring outcomes using polygenic scores (PGSs). However, there are concerns regarding potential biases due to confounding when mediating parental phenotypes are included.

**Methods:** Depression, educational attainment and height PGSs were generated for 2680 biological parent-offspring trios using genome-wide association study (GWAS) meta-analysis summary statistics in a large population study: Generation Scotland. Regression and pathway models were estimated incorporating PGSs for both parents and offspring to explore direct (offspring PGS) and genetic nurturing (parental PGS) effects on psychological distress, educational attainment and height. Genetic nurturing via parental phenotypes were incorporated into the models. To explore sources of bias we conducted simulation analyses of 10,000 trios using combinations of PGS predictive accuracy and accounted variance.

**Results:** Models incorporating both offspring and parental PGSs suggested positive parental genetic nurturing effects on offspring educational attainment, but not psychological distress or height. In contrast, models additionally incorporating parental phenotypic information suggested positive parent phenotype mediated genetic nurturing effects were at play for all phenotypes explored as well as negative residual genetic nurturing effects for height. 10,000 parent-offspring trio effects (without genetic nurturing effects) were simulated. Simulations demonstrated that models incorporating parent and offspring PGSs resulted in genetic nurturing effects that were unbiased. However, adding parental phenotypes as mediating variables results in biased positive estimates of parent phenotype mediated genetic nurturing effects and negative estimates of residual genetic nurturing effects. Biased effects increased in magnitude as PGS accuracy and accounted variance decreased. These biases were only eliminated when PGSs were simulated to capture the entirety of trait genetic variance.

**Conclusion:** Results suggest that in the absence of PGSs that capture all genetic variance, parental phenotypes act as colliders in the same way as heritable environments. Relatively simple models combining parental and offspring PGSs can be used to detect genetic nurturing effects in complex traits. However, our findings suggest alternative methods should be utilised when aiming to identify mediating phenotypes and potentially modifiable parental nurturing effects.

## INTRODUCTION

The standard quantitative genetic model partitions phenotypes into genetic and environmental effects (Falconer & Mackay 1996). However, evidence of gene environment correlations (rGE) i.e. where an individual’s genotype for a trait is also correlated with environmental influences, highlight that these effects may not be independent of one another (Gage et al, 2016). One way that genes and environments become correlated is through parental genetic nurturing effects. Parents can have a direct genetic effect by passing on half of their genome to form the offspring’s own genome, and a genetic nurturing effect by shaping the offspring’s rearing environment (Kong et al, 2018) ***(Figure 1)***.

**Figure 1.**
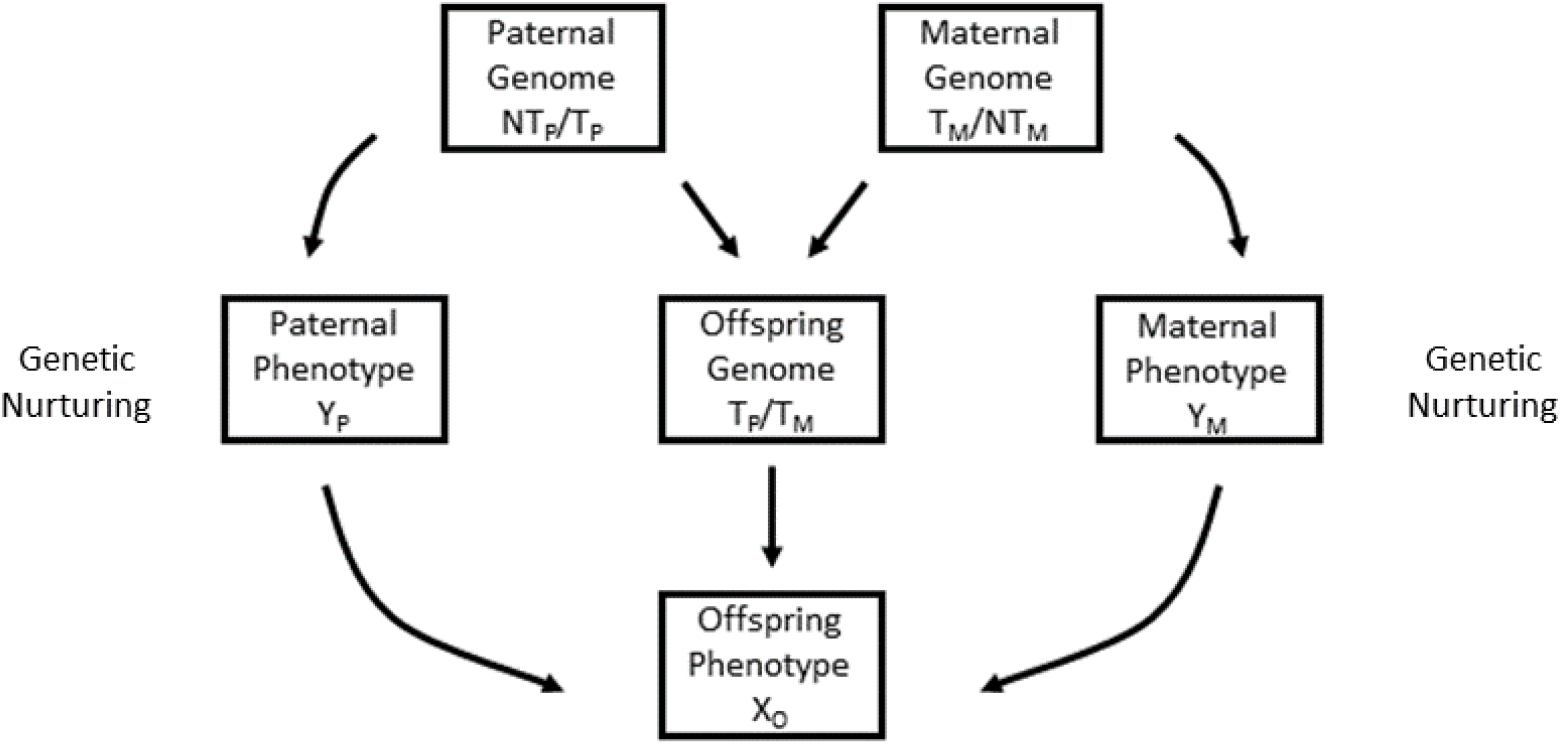
Direct genetic and genetic nurturing effects. Figure and legend adapted from Kong and colleagues (2018). T_P_ and T_M_ denote, respectively, the alleles transmitted from the father and mother. NT_P_ and NT_M_ denote the paternal and maternal alleles that are not transmitted. The paths show that the transmitted alleles can influence the phenotype of the offspring, X_O_, through a direct path. The paths also show that both transmitted and non-transmitted alleles can influence the parent phenotypes, Y_P_ and Y_M_, through which a genetic nurturing effect on the offspring phenotype, X_O,_ is observed. Whilst X is an individual trait of interest, Y may include a range of phenotypes that is not completely known.

Recently, the use and accuracy of polygenic scores (PGSs), derived from a weighted additive sum of trait associated alleles, has increased substantially over the last decade. Methods employing PGSs have suggested genetic nurturing effects in educational attainment by showing that scores were twice as predictive within non-adopted individuals in comparison to adopted individuals (Cheesman et al, 2020) as well as adoptive parental PGSs showing significant associations with adopted offspring education outcomes (Domingue & Fletcher, 2020).

Studies exploring trio PGSs, whereby direct genetic effects are controlled for by using offspring PGSs, provide evidence for potential genetic nurturing effects in educational attainment (Morris et al, 2020). Moreover, the use of phased family data has allowed for the partition and calculation of PGSs for alleles transmitted and not transmitted from parent to offspring (Kong et al, 2008). Findings using the non-transmitted PGSs demonstrate that genetic nurturing effects are present, albeit much smaller than direct genetic effects (Bates et al, 2018; Kong et al, 2018).

The estimation of genetic nurturing effects may provide a translational advantage by identifying parental behaviours or environmental factors that influence offspring phenotypes. This information has the potential to provide actionable findings or preventative treatment or policy. This has resulted in a steady accumulation of studies utilising PGSs within mediation analyses to explore particular environmental factors through which genetic effects may influence a given trait or disorder (Avinun, 2019; Avinun & Hariri, 2019; Shao et al, 2021; Stephan et al, 2020; Warrier & Baron-Cohen, 2019).

However, limitations associated with PGS predictive validity (i.e. accuracy of allele effect sizes and low numbers of associated genetic variants) are still present and have resulted in scores capturing only a fraction of the phenotypic variance for complex human traits (Dudbridge, 2013; Mostafavi et al, 2020). This, coupled with the lack of independence between genetic and environmental factors poses a challenge for statistical modelling of environment mediated PGS-trait associations (Akimova et al, 2020; Pingault et al, 2019), suggesting the need to revise the validity of these mediation analyses utilising PGSs.

This methodological challenge is particularly apparent when exploring parent phenotypes and parental genetic nurturing effects, as parent and offspring phenotypes are inevitably influenced by direct parental genetic effects. Consequently, the seemingly environmental variable (i.e. parent phenotype) is both heritable and genetically correlated with the offspring phenotype.

We sought to demonstrate the methodological challenges introduced with the addition of parental phenotypes in statistical models exploring parental genetic nurturing effects. We conducted simulations without genetic nurturing effects to highlight sources of genetic confounding and other potential biases that impact estimates of genetic nurturing utilising PGSs and parental phenotypes.

Here we focused on three different complex phenotypes: educational attainment, psychological distress and height. These phenotypes differ substantially in heritability estimates and PGS predictive validity. We utilised up to 2680 biological parent-offspring trios within the Generation Scotland sample. We used regression and pathway models to explore parental direct (offspring PGSs) and genetic nurturing effects (parental PGSs) on offspring phenotypes using trio PGSs to understand where misleading results can arise from mediation analysis.

## METHODOLOGY

### PARTICIPANTS

Participants were from Generation Scotland: The Scottish Family Health Study (GS), a family- and population-based study that recruited individuals from 2006 until 2011 (Smith et al, 2013). Currently GS has genetic, environmental and phenotypic data on over 20,000 individuals aged 18-99. To explore genetic nurturing effects from both parents to offspring, the study sample was limited to individuals with both biological parents genotyped (trios) resulting in 2680 parent-offspring trios. This sample includes 1596 singleton offspring, with the remaining offspring having at least one sibling within the sample. These sibling effects were controlled for by either including them as random effects into mixed linear models, where possible, or by including only one sibling in the analysis sample. The analysis samples were limited to trios with available phenotypic data on educational attainment (EA), psychological distress and height.

### PHENOTYPES

#### Educational Attainment

EA has relatively high heritability estimates of ∼66% (de Zeeuw et al, 2016; Schwabe et al, 2017) which is likely to yield greater statistical power in comparison to psychological distress. Furthermore, significant genetic nurturing associations with EA have been found within the literature (Bates et al, 2018; Cheesman et al, 2020; Kong et al, 2018; Morris et al, 2020). EA was available as a 1-10 point ordinal scale with higher values representing greater years of education.

#### Psychological Distress

Psychological distress is a moderately heritable trait, with twin and family study estimates ranging from 20-44% (Rijsdijk et al, 2003). The traits was assessed using the General Health Questionnaire (GHQ-28), a screening tool used to identify non-psychotic psychiatric disorders (Sterling, 2011). The GHQ-28 is well-validated and provides assessments of somatic symptoms, anxiety and insomnia, social dysfunction and severe depression (Goldberg & Hillier, 1979; Koeter, 1992). Each subsection is assessed using 7 behavioural items with a 4-point Likert scale (0-4) exploring the frequencies of experience: “not at all”, “no more than usual”, “rather more than usual”, “much more than usual”. This results in a minimum score of 0 and maximum score of 84.

#### Height

Similar to EA, height was explored as this is a trait with high heritability, with estimates showing approximately 85% of variation to be attributable to genetic differences (Silventoinen et al, 2003). Height has also yielded relatively well powered GWAS (Yengo et al, 2018) resulting in PGSs with substantial predictive validity i.e. accounting for a greater proportion of phenotypic variance in comparison to depression and educational attainment PGSs (Cesarini & Visscher, 2017). Height was measured in centimetres (cm).

Sample sizes for trios with available phenotypic data were 2,361, 2,315; 2,280; and 2,648 for EA, psychological distress, depression, and height, respectively. Information on sample demographics for each phenotype as well as information on age and sex distribution in the whole GS and trio samples are available in (Table S1).

#### GENOTYPES

Genetic data, single nucleotide polymorphisms (SNPs) were obtained from blood samples collected using standard operating procedures and genotyped using the IlluminaHumanOmniExpressExome-8v1.0 BeadChip and Infinium chemistry capturing 700,000 genome-wide SNPs and 250,000 exome SNPs (Kerr 2013). Quality control (QC) of genotyped SNPs consisted of the exclusion of SNPs with missingness > 2% and a Hardy Weinberg Equilibrium (HWE) p-value of ≤⍰1⍰×⍰10^−6^. SNPs with minor allele frequency (MAF) <0.01 and individuals with >3% missing genotypes were excluded from analyses. After QC procedures 561,125 SNPs were included in analyses.

#### POLYGENIC SCORES (PGSs)

EA, depression, and height PGSs for GS participants were created using allelic weight information from separate large GWAS meta-analyses ***(Table 2)***. To avoid over-fitting, GS participants were removed from these GWAS meta-analyses. The publicly accessible version of the EA GWAS was utilised (23&Me data within the GWAS meta-analyses were removed).

**Table 2.** Generation Scotland Polygenic Score Training Data

| Polygenic Score (PGS) | GWAS | GWAS Sample Size | Phenotype |
| --- | --- | --- | --- |
| Depression | (Howard et al, 2019) | 807,533 | Psychological Distress / Depression |
| Educational Attainment | (Lee et al, 2018) | 750,403 | Educational Attainment |
| Height | (Yengo et al, 2018) | 698,529 | Height |

*Note.* GWAS, Genome-wide Association Study

PRSice2 (Choi et al, 2020) with the default clumping (LD r2 < 0.1 across a 250 kb window) algorithm was used to compute the PGSs. To avoid biases that may arise in the clumping of SNPs from family-based data (GS data), an external reference panel (1000G linkage disequilibrium (LD) reference panel) was utilised.

Multiple PGSs were created using SNPs with varying trait association p-value thresholds (ranging from p < 5×10^−8^ to p <1). Identifying a single threshold that (a) results in PGSs that are not overfit to GS data and (b) is optimal for each phenotype is extremely difficult, especially as the traits being explored are substantially different to one another. To circumvent these potential limitations a principal components analysis was conducted on the different sets of PGSs for each phenotype separately and the ***first*** principal component/eigenvector (PC of PGSs) values were extracted. This method has been shown to produce scores that are not overfit and have increased predictive ability (Coombes et al, 2020).

The PC of PGSs were highly correlated with all different sets of PGSs, and each PGS uniformly loaded onto the PC (Table S2). PGS (including PC of PGSs)-phenotype associations were explored (Table S3) and the PC of PGSs values were found to be significantly associated with the relevant phenotype. The PC of PGSs was utilised as the trait PGS, from here onwards, this will be referred to as the PGS.

## STATISTICAL ANALYSES

### Generation Scotland Analyses

All statistical analyses were conducted in R (Team, 2020). Pearson’s correlation tests were conducted to explore trio PGS and phenotype correlations (S4). PGSs were standardised to have a mean of 0, and standard deviation of 1 and **all** variables were then pre-corrected for covariates; the first 20 genetic PCs of offspring and parents (to control for population stratification and cryptic relatedness), offspring and parental age and offspring sex. The residualised variables were used in subsequent regression and pathway models.

### Regression Analyses

Mixed effect multiple regression analyses were conducted (function ‘lmer’, package ‘lmerTest’) within R. Four separate models were explored and compared for the different phenotypes ***(Table 3)***:

**Table 3.** Regression Model Equations Model Equation

| Model | Equation |
| --- | --- |
| I | $oPheno \sim oPGS + Covariates + Sibling\ Effect$ |
| II | $oPheno \sim oPGS + mPGS + pPGS + Covariates + Sibling\ Effect$ |
| III | $oPheno \sim oPGS + mPheno + pPheno + Covariates + Sibling\ Effect$ |
| IV | $oPheno \sim oPGS + mPGS + pPGS + mPheno + pPheno + Covariates + Sibling\ Effect$ |

Note. Pheno, Phenotype; PGS, polygenic score; m/p/o, maternal, paternal, offspring respectively.

Model I aimed to capture direct genetic effects, i.e. the risk-associated effect of the offspring’s own genome (oPGS) on its phenotype. In contrast, model II aimed to capture both direct and parental genetic nurturing effects. If model II has a statistically better fit than model I, then any additional accounted variance can be attributed to parental genetic nurturing effects. Models III and IV aimed to provide additional insight using parental phenotypes as well as provide a comparison for the subsequent pathway models outlined below.

Offspring phenotypes (*oPheno*); EA, psychological distress, and height were included as dependent variables. Trio EA, depression and height PGSs (*oPGS, mPGS, pPGS*) and parental phenotypes (*mPheno, pPheno*) were included as predictors. Sibling effects were modelled as varying intercepts (random effects) for each sibling group to account for other sources of correlation between sibling phenotypes and avoid pseudoreplication. Sibling groups were identified as trio offspring who had the same parents.

Likelihood ratio tests were explored (r function ‘anova’) to compare the model fit of the four different models for each phenotype. Results include false discovery rate (FDR) adjustments for multiple testing (function ‘p.adjust’, q-value threshold <0.05) arising from the model comparisons (Table S6). The regression models were also explored using singleton, male and female offspring samples separately (Table S5).

### Pathway Analyses

Pathway analyses were conducted to explore the parental genetic and phenotypic effects on offspring traits. A key utility of pathway analyses is that the effect and proportion of variance captured by specific paths can be estimated. ***Figure 2*** presents a visualisation of the difference between multiple regression and pathway models.

**Figure 2.**
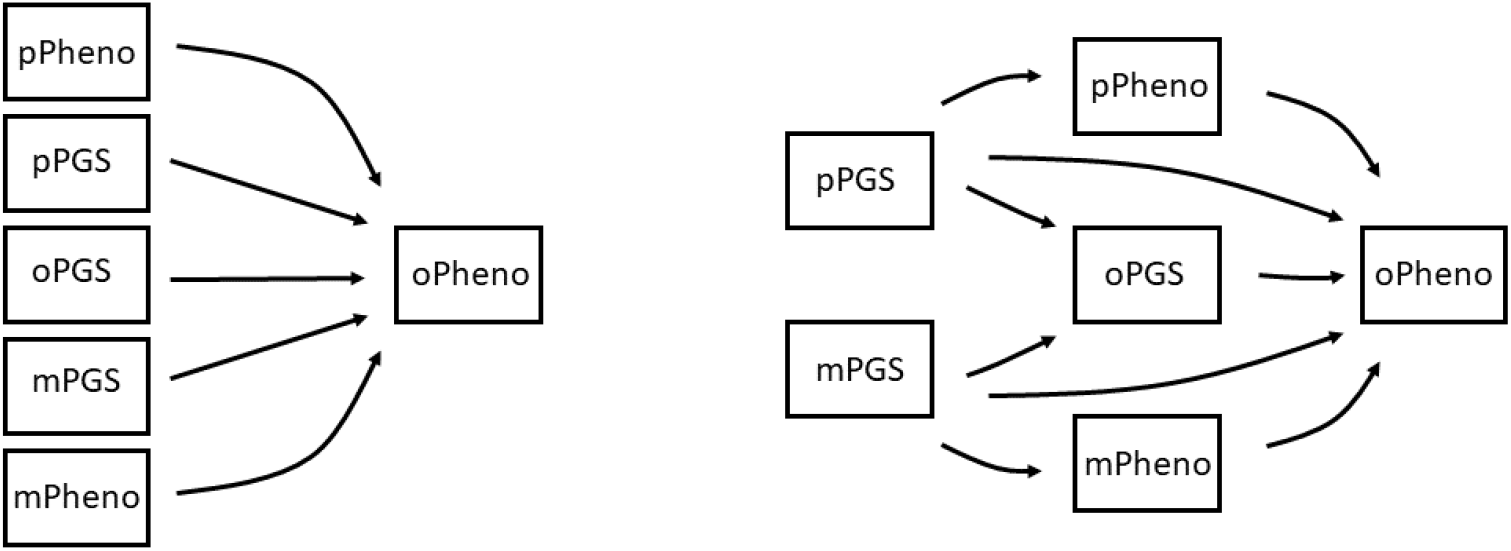
Visualisation of Multiple Regression and Pathway Models. PGS, polygenic score. Left, shows associations between the independent variables (offspring and parental PGSs and parental phenotypes) and the dependent variable (offspring phenotype). Right, shows associations between independent variables with the dependent variable whilst also capturing associations (mediation) between the independent variables forming pathways.

Pathway analyses were conducted using the software package Lavaan (Rosseel, 2012; 2018). Two pathway models with varying levels of complexity, labelled; simple and extended, were explored (***Figure 3)***. Similar to the regression analyses, offspring EA, psychological distress and height *(oPheno)* as well as trio EA, depression and height PGSs (*oPGS, mPGS, pPGS*) were included as predictors within the simple pathway analyses. The extended model builds upon this by further including parental phenotypes *(mPheno, pPheno)*. Associations within the simple and extended pathway models are outlined in ***Table 4***. The coefficients of associations between parental PGSs and offspring PGSs are fixed to 0.5, as parents and offspring are expected to share ∼50% of their genome. Pathway models were also conducted where all path coefficients were freely estimated, using singleton offspring (to eliminate sibling effects), only female and male offspring as well as for maternal and paternal duos separately (Table S7).

**Table 4.** Pathway Model Equations

| Pathway | Equation |
| --- | --- |
| *a* | $oPheno \sim oPGS$ |
| *b* | $oPheno \sim pPGS$ |
| *c* | $oPheno \sim mPGS$ |
| *d* | $pPheno \sim pPGS$ |
| *e* | $mPheno \sim mPGS$ |
| *f* | $oPheno \sim pPheno$ |
| *g* | $oPheno \sim mPheno$ |
| *h* | $oPGS \sim pPGS$ |
| *i* | $oPGS \sim mPGS$ |

*Abbreviations.* Pheno, Phenotype; PGS, polygenic score; m/p/o, maternal, paternal, offspring respectively.

**Figure 3.**
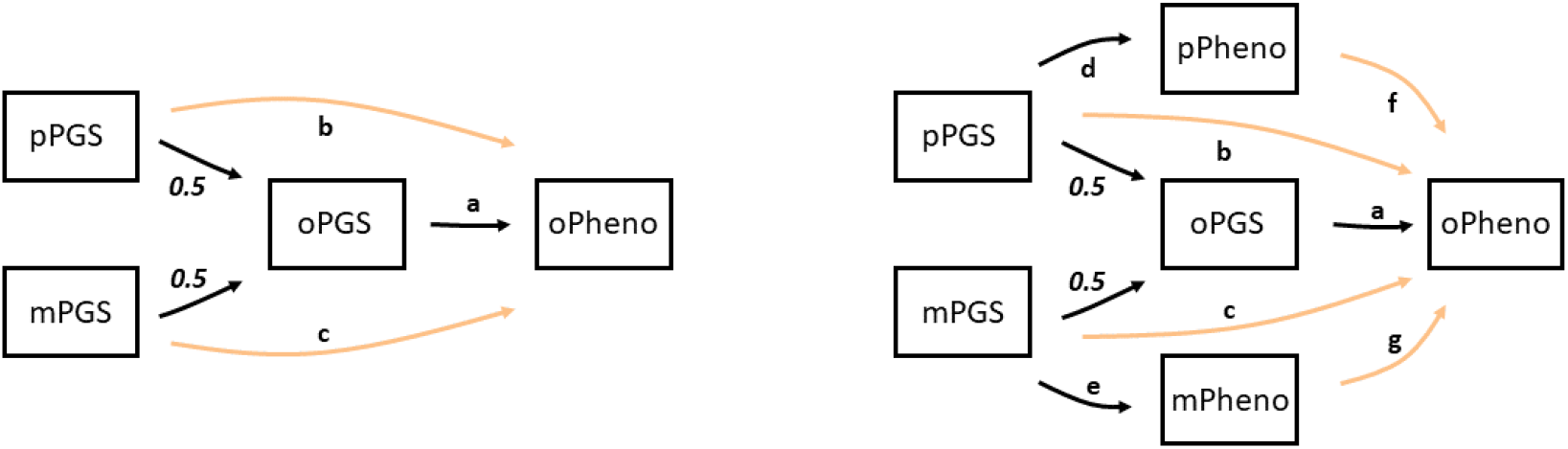
Simple and Extended Pathway Models. PGSs, polygenic scores; m, maternal; p, paternal; o, offspring. Left, Simple Model. Right, Extended Model. Associations between parental PGSs and offspring PGSs are fixed to 0.5 as parents and offspring share ∼50% of their genome. Paths 0.5*a represent direct genetic effects of parental PGSs on offspring phenotype. Paths b and c represent genetic nurturing effects of parental PGSs on offspring phenotype. Paths d*f and e*g represent the effect of parental PGS on offspring phenotype mediated by the parental phenotypes. Coloured pathways represent genetic nurturing effects.

### Simulation Analyses

Simulations were conducted to explore how results of the pathway models outlined above differ when using PGSs with varying levels of predictive ability. 10,000 trio members with respective genetic, phenotypic and PGS variables were simulated. These simulations did not include any genetic nurturing effects.

Simulated phenotypic variables had a variance of 1. Additive genetic variance was computed as the heritability multiplied by the phenotypic variance. Genetic variance was represented as two separate entities; tagged genetic variance, aiming to capture variance attributable to genotyped variants, and non-tagged genetic variance, aiming to capture variance attributable to non-genotyped variants. All remaining variance was attributed to environmental variance.

The parental genetic variables were constructed as the sum of tagged and non-tagged genetic counterparts. Tagged genetic variable values were simulated from a normal distribution with a mean of zero, and variance equal to the tagged genetic variance. Non-tagged genetic variable values were simulated from a normal distribution with a mean of zero, and variance equal to the non-tagged genetic variance.

Similarly, the offspring genetic variables were constructed as the sum of tagged and non-tagged genetic counterparts. Offspring tagged and non-tagged genetic variables were simulated as the average of the summed respective tagged and non-tagged maternal and paternal genetic variables, with the addition of a respective tagged and non-tagged segregation variable aiming to capture variability that occurs from random segregation of genes observed during meiosis (Yanowitz, 2010). The tagged and non-tagged segregation variable values were simulated from a normal distribution with a mean of zero, and variance equal to half the tagged and non-tagged genetic variance, respectively.

Separate environmental variables were constructed for each member of the trio. These variables were simulated from a normal distribution with a mean of zero, and a variance equal to the environmental variance.

The phenotypic variables for each member of the trio were then constructed as the sum of the respective trio member’s genetic and environmental variable values. PGS variables were derived from the trio member’s respective tagged genetic component. Different scenarios, with 15 replications each were conducted with a range of plausible trait heritabilities (0.3, 0.6, 0.9) and proportion of tagged genetic variances (0.2, 0.6, 1) for the findings to have broad applicability to the study of a wide variety of traits/complex trait architectures.

## RESULTS

### Generation Scotland Results

Offspring PGSs are correlated ∼0.5 with parental PGSs. Furthermore, weak significant associations for educational attainment (EA) and height PGS correlations are observed between the parents, consistent with evidence suggesting the presence of assortative mating (Conley et al, 2016; Hugh-Jones et al, 2016). Similarly, strong evidence of correlations between all trio members’ phenotypes was observed (Table S4).

The results of the four different regression models exploring direct and genetic nurturing effects are listed in Table S5. Log-likelihood comparisons of models I and II sought to identify genetic nurturing effects captured by parental PGSs. Results suggest that model II (including parental PGSs) accounts for greater variance and is significantly better than model I for EA only (**Table 5**). For all model comparison results see Table S6).

**Table 5.**
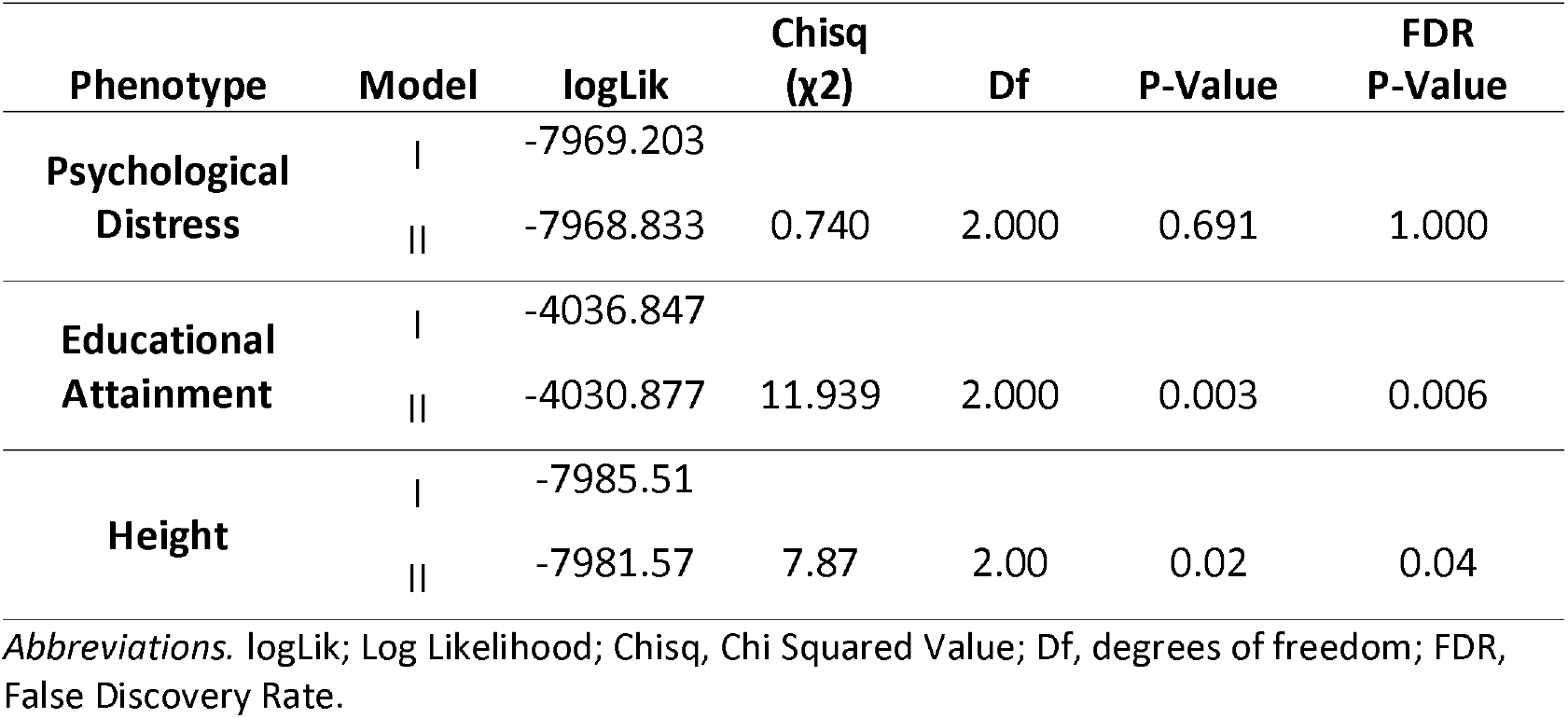
Regression Model I and II Log-Likelihood Comparisons

Notable results include negative and highly significant *β* coefficient estimates for parental height PGSs when included in models alongside parental height phenotypes (model IV). Interestingly, these estimates are positive and non-significant when included in models without parental phenotypes (model II).

Path *β* coefficient estimates, standard errors and p-values from the simple and extended pathway analyses are presented in **Figure 4**.

**Figure 4.**
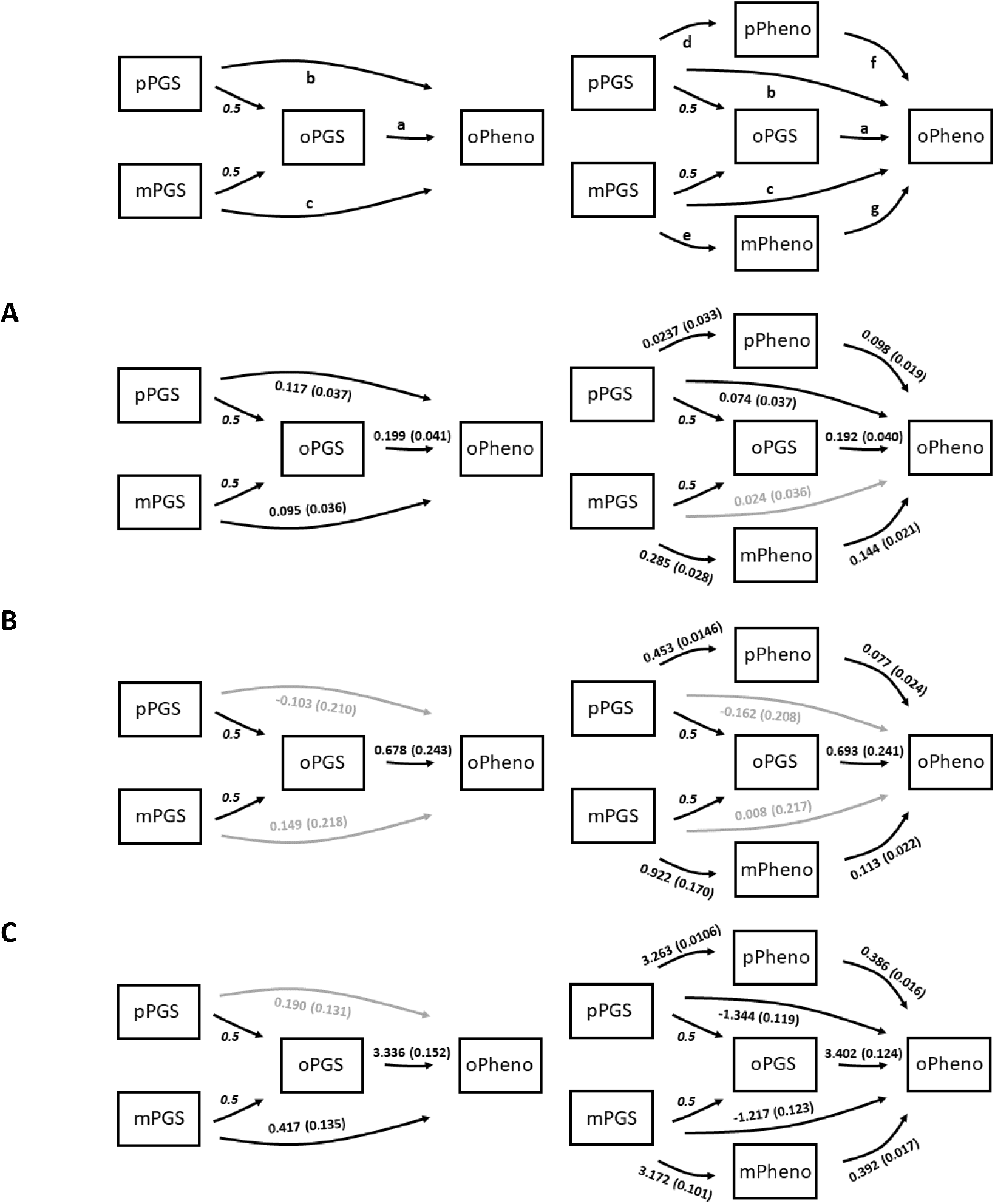
Simple and Extended Pathway Models Results. Left, simple; right, extended pathway models. Paths between parent PGSs and offspring phenotypes via offspring PGSs represent direct genetic effects of parental PGSs on offspring phenotype. The coefficients paths between parent PGSs and offspring PGSs are fixed to 0.5. Paths b and c represent genetic nurturing effects of parental PGSs on offspring phenotype. Paths *d*f* and *e*g* represent the effect of parental PGS on offspring phenotype mediated by the parental phenotypes. Panels A, B, C present path coefficients and standard errors for educational attainment, psychological distress and height, respectively. Grey paths represent associations that are *not* statistically significant (p>0.05).

The simple model results for EA suggest clear evidence of both paternal and maternal genetic nurturing effects on offspring educational attainment (paths *b* and *c*, =0.117, P<0.001, =0.095, P<0.001 respectively) which support and build upon the findings of the regression model comparisons. Paths representing the association between the trio PGSs and their respective phenotypes in the extended model, are quite different to each other, with the parental PGS and parental phenotypic associations being almost ∼2 fold greater than the offspring PGS and offspring phenotypic association (paths a, d and e; *β*=0.192, P<0.001; *β*=0.237, P<0.001; *β*=0.285, P<0.001, respectively). Furthermore, the inclusion of parental phenotypes within the extended model seem to result in a decrease in paternal genetic nurturing effect (path b, from *β*=0.117, P<0.001 to *β*=0.074, P<0.001) and the maternal genetic nurturing effect becoming redundant (path c) i.e. non-significant.

The simple and extended models show no evidence of genetic nurturing effects (panel A, paths b and c) on psychological distress. Notably, the extended model paths d (*β*=0.453, P<0.05) and e (*β*=0.992, P<0.001), exploring parental depression PGS and parental psychological distress associations, seem to be quite different, with maternal associations being ∼2 fold greater than paternal associations.

In contrast, associations between parental and offspring psychological distress from extended models (paths f and g, *β*=0.077, P<0.05 and *β*=0.113, P<0.001) suggests parental genetic nurturing effects mediated by parental psychological distress are at play. In fact, this pattern of mediated parental genetic nurturing effects are/were observed for all phenotypes.

The simple pathway model results suggest a small positive maternal genetic nurturing effect on offspring height (*β*=0.417, P<0.05). Extended model results show the parental genetic nurturing effect (paths b and c) become negative and highly significant (*β*=-1.344, P<0.001; *β*= -1.217, P<0.001), which is contradictory to the results from the simple model.

Patterns of results, and estimates are similar when all coefficients are freely estimated, as well as when samples are limited to singletons, female offspring, male offspring, maternal duos and paternal duos (Table S7).

### Simulation Results

Regression and pathway analyses were conducted using simulated trio data. Each analysis was repeated 15 times. As no genetic nurturing effects were simulated, it is expected that paths *b, c, f* and *g* of the pathway models will be approximately 0 and non-significant when PGSs are adequately capturing genetic variance (**Figure 3**).

Figure 5. presents *β* coefficient boxplots of paternal genetic nurturing paths highlighted in **Figure 3**. This figure shows that when the PGSs are perfect measures of the genetic variance, i.e. tags the entirety of trait genetic variance, the estimated paternal genetic nurturing (paths *b* and *f*) coefficients are close to 0. In all other scenarios, extended model estimates for path *b* are always negative and estimates for *f* are always positive. These effects are exacerbated as trait heritability increases and variance captured by PGSs decreases. Results from regression models, and maternal genetic nurturing paths show similar *β* coefficient estimate patterns (Figure S1-3).

## DISCUSSION

The present study utilises polygenic scores (PGSs) informed by large international GWAS consortia and 2680 Generation Scotland trios. Regression and pathway models using available GS data found contradicting results. Models with only trio PGSs (regression models II and simple pathway models) showed evidence of genetic nurturing effects for educational attainment (EA) only. However, the addition of parental phenotypes into models (regression models IV and extended pathway models) showed parental phenotype mediated genetic nurturing effects for *all* phenotypes explored. Moreover, highly significant and negative residual parental genetic nurturing effects (i.e. not mediated by parental phenotypes) were observed for height.

**Figure 5.**
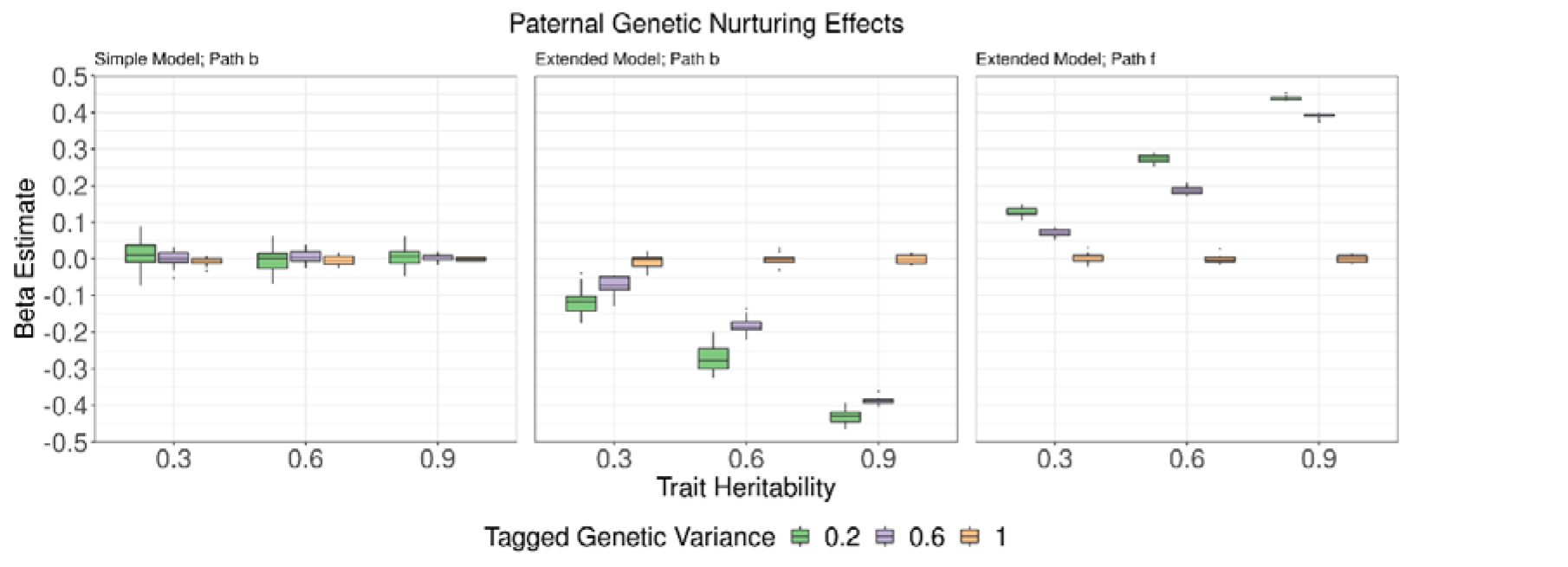
Paternal Genetic Nurturing Effects. Boxplots represent *β* coefficients from 15 replication analyses. Colours present simulated polygenic scores (PGS) capturing varying levels of trait genetic variance. X-axis presents varying levels of simulated trait heritability; y-axis presents the beta coefficients of genetic nurturing effects. The left and middle plots present boxplots representing *β* coefficient estimates of path b; the association between offspring phenotype and paternal PGS from the simple and extended pathway models respectively. The right plot presents boxplots representing *β* coefficient estimates of path f from the extended pathway model; the association between offspring phenotype and paternal phenotype.h

Analyses of simulated genetic nurturing models showed that utilising PGSs that did not capture the total genetic variance, whilst including both parental PGSs and parental phenotypes as predictors, resulted in biased estimates of nurturing effects. Parental phenotypes were always positively associated with the offspring phenotypes (paths f and g), resulting in what seemed to be significant mediated parental genetic nurturing effects, despite expectations of null associations (as genetic nurturing effects were not simulated). Significant associations will arise between parental phenotypes and offspring phenotypes due to shared genetic variance that is not adequately accounted for by the PGSs within the models; leading to the potentially erroneous interpretation of significant mediated parental genetic nurturing effects. Biased estimates were also observed for the residual parental genetic nurturing effects. These estimates were negative, again despite expectations of null associations. Results further highlight that these upward and downward biases were exacerbated, as PGSs became poorer measures of genetic variance and trait heritability increased. These biases were only eliminated when PGSs were simulated to capture the entirety of trait genetic variance. Thus, in the absence of PGSs that capture all of the genetic variance, parental phenotypes act as colliders in the same way as heritable environments (Akimova et al, 2020).

Results using simulated data highlight the confounded effects in the models including parental phenotypes that are not necessarily captured by observable data. Confounded effects can explain the discrepancy observed between simple and extended model results using GS data. In fact, GS height extended model results were highly comparable to the pattern of results using simulated data, which may result from the lack of/negligible genetic nurturing effects as suggested by the GS height simple model results. In contrast, results from analyses using GS EA and psychological distress being dissimilar to that observed from analyses using simulated data, may suggest genuine genetic nurturing effects at play.

The extended pathway models using GS data, also included discrepancies observed in the association effect sizes between respective trio PGS and phenotype associations. Maternal PGS-phenotype associations were much larger than that observed for fathers when exploring psychological distress, suggesting potential sex interaction effects, which may require further investigation. Similarly, parental PGS-phenotype associations were larger than observed offspring PGS-phenotype associations for EA. This may be attributable to the fact that genetic nurturing effects play a significant role in EA, and the absence of grandparent PGSs within the models may result in parental PGS-phenotype associations encompassing additional genetic nurturing effects. These effects were accounted for within offspring PGS-phenotype associations, as parent data is included in the models.

Overall, results suggest that the regression and simple pathway models exploring offspring PGS (direct genetic effects) and additional parental PGS (genetic nurturing effects) associations with offspring phenotypes, pose a straightforward and unbiased method to explore genetic nurturing effects.

Log-likelihood comparisons of regression models (I and II) and simple pathway model results also suggested evidence of parental genetic nurturing effects for EA, but not other explored phenotypes. The EA regression model with all trio member’s PGS included accounted for greater variance and showed a significantly better fit in comparison to the model using only the offspring PGS. This supports much of the findings from the literature suggesting genetic nurturing effects are detectable using current EA PGSs (Bates et al, 2018; Kong et al, 2018).

Whilst these findings highlight future research directions, the biased results using simulated data clearly show that the extended models are confounded. It is evident that without essentially perfect PGSs, accurate quantification of genetic nurturing effects from extended models is not possible.

Future work can include the adaptation of these models to utilise data on different traits between parents and offspring e.g. parent EA PGS and offspring depression PGS to explore genetic nurturing effects of parental EA on offspring depression. It is important to keep in mind differences in GWAS power, and thus, discrepant predictive validity of these PGSs may result in further biases. Ideally, additional simulation analyses should be conducted to explore the impact of differences in parent and offspring PGS power on results.

As with all research designs, some limitations should be considered when interpreting the results presented. Here trio data associations were simulated using random sampling from pre-defined variances, as opposed to simulating individual genotypes. Whilst this was adequate in highlighting the biases that may arise within parental genetic nurturing models proposed; it is a simplistic representation of the effects shared between biological parent-offspring trios. For example, genetic nurturing effects may be at play via both parental transmitted and non-transmitted alleles. Thus, potential differences in how biases may arise within these specific paths were not observed. Moreover, inaccurate effect sizes obtained from GWAS summary statistics may also contribute to noise in PGSs, which is another level of complexity not modelled here. Additionally, it is important to note that shared environmental effects were not modelled in analyses here, however, may also drive significant associations and confound effects within the extended models. Identifying and modelling all relevant environmental effects is likely not possible, and thus, these model results may always be biased to some extent.

Another limitation not formally explored is the impact of assortative mating within these models. In fact, evidence of assortative mating has been observed for many complex behavioural traits, including phenotypes explored within this study (Hugh-Jones et al, 2016; Mathews & Reus, 2001; Stulp et al, 2017). Strong evidence of assortative mating, yet limited evidence of genetic nurturing effects for height has been shown within the literature (Stulp et al, 2017). Here, unbiased results of GS height analyses showed no evidence of genetic nurturing, which suggests that any potential bias arising from assortative mating effects are likely to be very small.

This study demonstrates that pathway models are a simple and useful method to explore and quantify genetic nurturing effects using offspring phenotypic and trio PGS data. However, it is important to be aware of biases which arise when including parental phenotype mediated genetic nurturing paths. The improvement of PGSs will likely fall short for the level required for clear interpretation of these models. These results suggest alternative methods should be utilised when exploring factors through which genetic nurturing effects may be at play.

## DATA & CODE AVAILABILITY

Data is available to researchers upon application to http://generationscotland.org. Code is available from https://github.com/melisachuong/rGE_Depression.

